# A haplotype-resolved reference genome for *Eucalyptus grandis*

**DOI:** 10.1101/2025.02.07.637144

**Authors:** Anneri Lötter, Tomas Bruna, Tuan A. Duong, Kerrie Barry, Anna Lipzen, Chris Daum, Yuko Yoshinaga, Jane Grimwood, Jerry W. Jenkins, Jayson Talag, Justin Borevitz, John T. Lovell, Jeremy Schmutz, Jill L. Wegrzyn, Alexander A. Myburg

## Abstract

*E. grandis* is a hardwood tree used worldwide as pure species or hybrid partner to breed fast-growing plantation forestry crops that serve as feedstocks of timber and lignocellulosic biomass for pulp, paper, biomaterials and biorefinery products. The current v2.0 genome reference for the species (Bartholome et al., 2015; Myburg et al., 2014) served as the first reference for the genus and has helped drive the development of molecular breeding tools for eucalypts. Using PacBio HiFi long reads and Omni-C proximity ligation sequencing, we produced an improved, haplotype phased assembly (v4.0) for TAG0014, an early-generation selection of *E. grandis.* The two haplotypes are 571 Mbp (HAP1) and 552 Mbp (HAP2) in size and consist of 37 and 46 contigs scaffolded onto 11 chromosomes (contig N50 of 28.9 and 16.7 Mbp), respectively. These haplotype assemblies are 70 to 90 Mbp smaller than the diploid v2.0 assembly but capture all except one of the 22 telomeres, suggesting that substantial redundant sequence was included in the previous assembly. A total of 35,929 (HAP1) and 35,583 (HAP2) gene models were annotated, of which 438 and 472 contain long introns (>10 kbp) in gene models previously (v2.0) identified as multiple smaller genes. These and other improvements have increased gene annotation completeness levels from 93.8% to 99.4% in the v4.0 assembly. We found that 6,493 and 6,346 genes are within tandem duplicate arrays (HAP1 and HAP2, respectively, 18.4% and 17.8% of the total) and >43.8% of the haplotype assemblies consists of repeat elements. Analysis of synteny between the haplotypes and the *E. grandis* v2.0 reference genome revealed extensive regions of collinearity, but also some major rearrangements, and provided a preview of population and pan-genome variation in the species.

**Paper summary:** We assembled a haplotype-phased genome for *Eucalyptus grandis* that will serve as reference for the most widely planted hardwood crop globally. It includes more than 430 new gene models with long introns and has 6% higher annotation completeness. The phased assembly provides a more accurate look at genome variation at DNA and transcript level and will better support future studies of genome structure and function. The improved assembly contains more tandem duplicate genes compared to the previous unphased reference. Finally, major genomic rearrangements between the two phased genomes provide a preview of pangenome and structural variation in *E. grandis*.

## Introduction

The eucalypts are a group of woody plants with more than 900 species (Brooker, 2000) including a number of fast-growing forestry species that are being domesticated as timber and woody biomass feedstocks for a bio-based economy. Together with natural stands and restoration forestry efforts (Bennett, 2016; Brancalion et al., 2019; Jordan et al., 2016; Supple et al., 2018), eucalypt plantations can contribute to global carbon drawdown (Bar-On et al., 2018). *Eucalyptus grandis* is the model species for the genus, representing the most widely planted hardwood crop in the world (more than 20 million ha, Iglesias-Trabado & Wilstermann, 2008; Myburg et al., 2014). *E. grandis* is a highly outbred and heterozygous species (Myburg et al., 2014), with a natural habitat spanning much of the east coast of Australia (−16.2494, 145.3213 to −32.9305, 151.7541) and multiple climatic conditions (Mostert-O’Neill et al., 2021). Due to the commercial importance of *E. grandis* (Harwood, 2011) and its position as a genetic model for the genus, numerous genomic (Bartholome et al., 2015; Myburg et al., 2014; Silva-Junior et al., 2015) and transcriptomic (Carocha et al., 2015; Hussey, Saidi, et al., 2015; Mangwanda et al., 2015; Mizrachi et al., 2010; Mizrachi et al., 2017; Oates et al., 2015; Soler et al., 2015; Vining et al., 2015) resources have been generated which have greatly aided understanding of the population genetics (Mostert-O’Neill et al., 2021) and biology (Hussey, Saidi, et al., 2015; Mangwanda et al., 2015; Mizrachi et al., 2017; Oates et al., 2015; Soler et al., 2015; Wierzbicki et al., 2019) of *E. grandis,* while promoting the development of molecular breeding tools in the genus (Ballesta et al., 2019; Candotti et al., 2023; Mostert-O’Neill et al., 2022; Silva-Junior et al., 2015; Silva-Junior & Grattapaglia, 2015; Telfer et al., 2015). These resources depend on the current reference genome, which was assembled from whole-genome Sanger and BAC-end sequences of a partially inbred genotype (BRASUZ1) and likely represents a collapsed assembly in the remaining heterozygous regions of the BRASUZ1 genome, rather than a haplotype phased assembly, which is rapidly becoming the standard in outbred organisms. Despite having proven a very good reference for the applications described above, the v2.0 assembly (Myburg et al., 2014) and the improved v2.1 assembly (Bartholome et al., 2015) would not have provided robust assessment of genome sequence and structural diversity in the species.

Long-read sequencing technologies promise improved, phased genome assemblies in outbred organisms, along with full-length transcript sequences that result in higher quality annotations. Such phased references allow identification of haplotype-specific variants (including structural variants, SVs) and the subsequent association of such variants with traits of interest (Cai et al., 2022; Healey et al., 2024; Scarlett et al., 2023; Shang et al., 2022; Tang et al., 2021; Vourlaki et al., 2022; Wang et al., 2023). Large SVs contribute to the genetic diversity of the species (pangenome variation) and lead to genome and 3D architectural changes that can alter gene expression regulation (Alonge et al., 2020; Kang et al., 2023; Li et al., 2024; Ni et al., 2023; Ni & Tian, 2023; Zhang et al., 2024). Ultimately, at individual level, such regulatory variation may contribute to phenotypic plasticity, the ability to survive diverse environmental challenges.

Here we report a haplotype phased genome reference for *E. grandis*, which is highly contiguous and nearly complete. This improved reference provides an important resource for studies focusing on genome sequence and structural evolution in the species compared to earlier studies that relied on sparse SNP markers alone (Mostert-O’Neill et al., 2021; Mostert-O’Neill et al., 2022). In addition, as an early generation commercial clone, the TAG0014 reference will be useful for studies of the genetic consequences of early domestication and breeding. Together with other recent phased genome assemblies in the genus (Ferguson, Bar-Ness, et al., 2024; Ferguson, Jones, Murray, Andrew, Schwessinger, & Borevitz, 2024; Ferguson, Jones, Murray, Andrew, Schwessinger, Bothwell, et al., 2024) the phased *E. grandis* (TAG0014, v4.0) assembly and improved gene annotation will also improve understanding of gene function and evolution, and support tree biotechnology efforts aimed at improving tree growth, development and resilience to climate change and associated biotic challenges.

## Methods and Materials

### Genome sequencing and haplotype-phased assembly

We sequenced an early selection (first generation from wild, unimproved material) of *E. grandis* which has been clonally propagated since the 1990s but is no longer used as a commercial clone. The availability of ample clonally replicated trees (Mondi South Africa) allowed us to perform whole-tree collection of developing tissues for genome and transcriptome analysis. Due to its high susceptibility to pathogens in subtropical regions (Wingfield et al., 2008), *E. grandis* is often used as a hybrid partner, typically with *E. urophylla,* for clonal plantations of F1 hybrid varieties in these regions. As a clone that was derived from early breeding trials involving seed from unimproved material from several provenances, TAG0014 is possibly an inter-provenance cross of *E. grandis* with higher heterozygosity than individuals from native provenance stands. It was selected to be the new reference due to its early generation status, availability as a clonal genotype for molecular studies and because of the numerous datasets already generated for this clone (Hussey et al., 2017; Hussey, Mizrachi, et al., 2015; Vining et al., 2015).

The TAG0014 genome was sequenced with a range of sequencing technologies including PacBio HiFi (Pacific Biosciences Inc., CA, USA), chromosome conformation capture with Omni-C (Dovetail Genomics, CA, USA) and Illumina short-read sequencing data. PacBio Iso-Seq and Illumina RNA-seq data was generated for genome annotation. High molecular weight (HMW) DNA was isolated from frozen leaf tissue at the Arizona Genomics Institute (AGI, Tucson, Arizona, USA). PacBio and Illumina sequence reads were generated at the HudsonAlpha Institute for Biotechnology (HA) in Huntsville, Alabama and at the Department of Energy Joint Genome Institute (DOE-JGI) in Berkeley, California. An Illumina library (400 bp insert) was generated and sequenced on the NovoSeq 6000 platform to obtain 2×150 bp paired-end reads to a coverage of 64.4x along with a 2×150 nt Dovetail Omni-C library (**Supplementary Figure 1**). PacBio sequence reads were generated using the Sequel IIe platform at HA.

Error-corrected PacBio reads (41.92 Gbp, 65.50x coverage and read N50 of 17.74 kbp) were assembled into initial haplotype phased contigs using HiFiAsm+HiC (Cheng et al., 2021). The assemblies were polished with RACON (Vaser et al., 2017) and no misjoins were identified in the polished assemblies. This produced highly contiguous initial assemblies for both haplotypes. The assemblies were further scaffolded, ordered, and orientated using Omni-C proximity ligation contacts generated with the JUICER pipeline (Durand et al., 2016) and the *E. grandis* v2.0 genome. Contigs terminating in significant telomeric sequences were properly oriented in the assembly. For HAP1, a total of 13 joins resulted in the final assembly of 11 chromosomes (containing 99.991% of the assembled sequence) and one small unplaced scaffold of 50 kbp. Adjacent alternative haplotypes were identified on the joined contig set. Seven adjacent alternative haplotype (althap) regions were collapsed in the assembly using the longest common substring between the two haplotypes. Similarly, for HAP2, 40 joins were applied to obtain a final assembly of 11 chromosomes which contains 100% of the assembled sequence. A total of 16 adjacent althap regions were collapsed in the HAP2 assembly. The resulting assemblies were screened for contaminants using NCBI’s FCS-GX pipeline (Astashyn et al., 2024). The mitochondrial and chloroplast genomes were assembled using OatK (https://github.com/c-zhou/oatk). Additional scaffolds were classified as redundant (63/100 HAP1/HAP2 scaffolds composed of >=95% 24mers >2x in all scaffolds, 8.9/13.9 Mbp), repetitive (39/21 <=250 kbp scaffolds composed of >=95% 24mers >4x in >=5 Mbp scaffolds, 3.9/1.5 Mbp) or organellar (assembled with OatK before genome assembly, 456.9 kbp mitochondrial genome, 160.2 kbp chloroplast).

HifiAsm may occasionally misphase small regions within the chromosomes (Healey et al., 2024). Misphased regions in the TAG0014 haplotypes (v3.0) were identified through alignment of Omni-C data to the combined HAP1/HAP2 chromosomes and manual identification of the misphased regions in the contact map. The corrected haplotype-specific phased assemblies were the final assemblies selected as haplogenome references (v4.0). The genome assemblies were screened for PacBio linkers, and 14/13 instances of linkers were found in HAP1/HAP2. When comparing haplotypes, we noted regions on Chr01 HAP2 that were absent in HAP1. Closer inspection revealed that the HAP2 region had two-fold higher depth than the surrounding regions suggesting that it is likely a collapsed homozygous copy. In the v4.0 TAG0014 release, the entire homozygous region identified by a doubling of CCS depth was duplicated in HAP1.

Homozygous SNPs and INDELs were corrected in both haplotypes using ∼60.4x Illumina reads. A total of 68/118 (HAP1/HAP2) SNPs and 3,753/4,048 INDELs were corrected for HAP1/HAP2. Heterozygous SNPs and INDELs were corrected using 47.75x CCS data, fixing 90/44 SNPs and 654/627 INDELs for HAP1/HAP2, respectively.

Genome assembly statistics were generated with QUAST v5.2.0 (Gurevich et al., 2013; Mikheenko et al., 2018) and completeness was assessed with BUSCO v5.4.5 (Manni et al., 2021; Seppey et al., 2019; Simao et al., 2015) using the embryophyta_odb10 database. Genome contiguity of the TAG0014 phased (v4.0) and BRASUZ1 (v2.0) diploid genome reference was visualised with GENESPACE (Lovell et al., 2022).

Completeness of the euchromatic portion of the genome assembly was assessed by aligning 34,121 annotated primary transcripts from the *E. grandis* v2.0 annotated primary transcripts to the v4.0 HAP1 and HAP2 release. The completeness analysis aims to obtain a measure of completeness of the assembly, rather than a comprehensive examination of the gene space. We retained genes that aligned at greater than 90% identity and 85% coverage. We found that 32,663/32,577 (95.73%/95.47%) of the previously annotated genes align to the HAP1/HAP2 v4.0 release. Of the remaining annotated genes, 1,068/1,171 (3.13%/3.43%) aligned at <50% coverage and 390/373 (1.14%/1.09%) of genes were not found in the HAP1/HAP2 v4.0 release.

### Genome annotation

Total RNA was extracted from 15 tissues representing different developmental stages and positions on the TAG0014 tree (**Supplementary Figure 1**) using the Plant/Fungi total RNA purification kit (Norgen Biotek Corp.) per the manufacturer’s protocol. RNA purity was assessed using spectroscopy on a NanoDrop ND1000 (Nano-Drop Technologies) and integrity was verified with an Agilent 2100 Bioanalyzer (Agilent Technologies, Santa Clara, CA, USA). RNA-Seq data was generated by the JGI (Berkeley, CA) for the 15 tissue libraries prepared using Illumina’s TruSeq Stranded mRNA HT sample prep kit using poly-A selection of the mRNAs. The libraries were sequenced on the NovaSeq S4 platform to produce 2×150 bp RNA-seq reads. Transcripts were assembled from ∼367M 2×150 bp stranded PE reads with PERTRAN (Shu et al., 2013), which conducts genome-guided transcriptome short read assembly via GSNAP (Wu & Nacu, 2010) and builds splice alignment graphs after alignment validation, realignment, and correction.

PacBio Iso-Seq full-length RNA transcript sequences were generated using the Sequel IIe platform for three tissue pools (developing cambium/woody, green/leaf and flower) generated using pools of the RNA extracted above. A genome-guided correction pipeline was used to correct and collapse 25.9 M PacBio Iso-Seq CCS reads. The pipeline aligns CCS reads to the genome with GMAP (Wu & Nacu, 2010), corrects small indels in splice junctions, and clusters alignments when all introns are the same or >= 95% overlap for single-exon alignments. This resulted in ∼915K/908K putative full-length transcripts for HAP1 and HAP2 respectively. Transcript assembly sets were combined into a final set of 364,350 HAP1 transcripts and 359,285 HAP2 transcripts with PASA (Haas et al., 2003).

A *de novo* repeat library was created for the TAG0014 HAP1 v3.0 genome draft using RepeatModeler2 (Flynn et al., 2020). To identify repeats with significant hits to protein-coding domains, the repeat library was functionally analysed with InterProScan (Jones et al., 2014) based on the Pfam (Mistry et al., 2021) and PANTHER (Mi et al., 2019) databases. Repeats with significant protein-coding domain hits were removed from the repeat library and the genome was soft-masked using RepeatMasker (Smit et al., 2013) with the resulting species-specific repeat library.

To determine putative gene loci, transcript assembly alignments and/or EXONERATE (Slater & Birney, 2005) alignments of proteins from *Corymbia citriodora*, *Arabidopsis thaliana*, *Glycine max*, *Fragaria vesca*, *Vitis vinifera*, *Liriodendron tulipifera*, *Punica granatum*, *Rhodamnia argentea*, *Syzygium oleosum*, *Brassica rapa*, *Citrus clementina*, *Gossypium raimondii*, *Populus trichocarpa*, *Oryza sativa*, *Beta vulgaris*, and Swiss-Prot release 2022_04 of eukaryote proteomes were generated using repeat-soft-masked *E. grandis* var. TAG0014 HAP1 and HAP2 v4.0 genomes, with up to 2 kbp bidirectional extension unless extending into another locus on the same strand. Gene models in each locus were predicted by homology-based predictors, FGENESH+ (Salamov & Solovyev, 2000), FGENESH_EST (similar to FGENESH+, but using EST to compute splice site and intron input instead of protein/translated ORF), EXONERATE, PASA assembly ORFs (in-house homology constrained ORF finder), and AUGUSTUS (Stanke et al., 2006) trained on the high confidence PASA assembly ORFs and with intron hints from short read alignments. The best-scored predictions for each locus were selected using multiple positive factors including EST and protein support, and one negative factor: overlap with repeats. The selected gene predictions were improved by adding UTRs, splicing correction and adding alternative transcripts with PASA.

To obtain a Cscore (the protein BLASTP score ratio to the mutual best hit BLASTP score) and protein coverage (the percentage of protein aligned to the best of homologs), PASA-improved gene model proteins were subject to protein homology analysis to the above-mentioned proteomes. PASA-improved transcripts were selected based on Cscore, protein coverage, EST coverage, and their CDS overlap with repeats. The transcripts were selected if their Cscore and protein coverage were >= 0.5 or if covered by ESTs. For gene models whose CDS overlapped with repeats by more than 20%, the Cscore had to be at least 0.9 and homology coverage at least 70% to be selected. The selected gene models were subject to Pfam analysis and gene models without strong transcriptome and homology support and whose proteins overlapped more than 30% with Pfam transposable element domains were removed. Incomplete gene models, low homology supported without full transcriptome-supported gene models, short single exon (< 300 bp CDS) without protein domains nor good expression, and repetitive gene models without strong homology support were manually filtered out.

Detailed gene metrics were obtained for the TAG0014 haplotypes and for the BRASUZ1 v2.0 genome with gFACs v1.1.2 (Caballero & Wegrzyn, 2019). Completeness of transcript and protein models was determined using BUSCO v5.4.5 and the embryophyta_odb10 database.

### Improvement of reference genome

The improvement of the v4.0 phased reference over that of the v2.0 draft was evaluated using the assembly and annotation completeness, number of gene models, assembly contiguity, genome feature presence, whether the assembly was phased and by looking for assembly gaps that have been closed. Telomere sequences were identified for all genomes using the (CCCGAAA)n and (CCCTAAA)n repeats with GENESPACE. Overall conservation of genome structure was evaluated through genome-wide, and gene-based synteny analyses.

Genome-wide synteny was assessed with SyRI (Goel et al., 2019). Pairwise combinations of reference genome assemblies were aligned using nucmer from the MUMmer4 toolbox (Marcais et al., 2018). Resulting alignments were filtered with nucmer (“—maxmatch –c 100 -b 500 −l 50”). The alignments were further filtered for alignment length (>100) and identity (>90), then used to identify local variants and structural rearrangements with SyRI v1.6.3 and visualised with plotsr v1.1.1 (Goel & Schneeberger, 2022).

The overall conservation of synteny of orthologous and homeologous gene regions was assessed using GENESPACE (Lovell et al., 2022). GENESPACE is based on protein similarity scores to construct synteny blocks with MCScanX (Wang et al., 2012) and Orthofinder v2.5.5 (Emms & Kelly, 2015, 2019). Syntenic blocks were used to identify pairwise peptide differences among the *E. grandis* genomes.

One of the main features of the BRASUZ1 reference genome was the high percentage of tandem duplicate arrays (12,570 genes in 3,185 clusters, Myburg et al., 2014). Since technological and methodological advances have been made since the original publication, we re-evaluated tandem duplicate gene content across all of the *E. grandis* reference genomes. Using the “combed.txt” output generated from GENESPACE, all orthogroups with more than two genes were considered as a tandem array and the number of genes and number of arrays were extracted using custom R scripts. The number of genes per array were also compared to find arrays where there are expansion/contractions relative to the current v2.0 genome reference.

### Population diversity analyses

The genetic relatedness of the TAG0014 genome to natural populations (provenances) of *E. grandis* was assessed using a 72K SNP Axiom 384-format array for *Eucalyptus* and *Corymbia* (Affymetrix Inc., Santa Clara, California, USA). Allelic intensity data were first assessed using Axiom Analysis Suite v5.2.0.65 to recluster genotypic classes as described by Silva-Junior et al., 2015. SNP data for natural population references was obtained from Mostert-O’Neill et al., 2021 and the SNP calls were converted to Axiom SNPs using a custom R script in RStudio v2023.12.1. The converted calls were imported into SNP & Variation Suite™ v8.9.1 (SVS8; Golden Helix, Inc., Bozeman, MT). High confidence, informative SNPs were retained by filtering for minor allele frequency >0.02, genotyped in at least 80% of samples and Hardy Weinberg Equilibrium P-value > 1e-5. Starting with 18,328 SNP markers that mapped to the v2.0 reference genome, 17,036 SNPs remained after filtering. To determine the genetic relatedness of the TAG0014 genome with trees representing the natural range of *E. grandis*, SNP data from Mostert-O’Neill et al., 2021 and the TAG0014 genome was used to perform a Principal Component Analysis (PCA, Price et al., 2006) in SVS8.

## Results

### Genome assembly and annotation

The *E. grandis* v2.0 reference had some limitations that prohibited its use to study larger SVs that contribute to pangenome variation in the species. First, due to the sequencing technology used and genome assembly capabilities available at the time, repeat regions were difficult to resolve. As a result, the current v2.0 genome reference was a fragmented assembly consisting of 21,856 contigs in 4,943 scaffolds (**Figure 1**, **Table 1**). Of the 640.4 Mbp v2.0 scaffolded reference, 7.4% (47.4 Mbp) is contained in gaps which may be the result of an overinflated genome size estimate (i.e. a forced genome size fit based on C-size estimates) and, to some extent, the retention of heterozygous genomic regions in the assembly. Recent studies suggest a smaller genome size when haplotype phased genome assemblies are generated (Lotter et al., 2023; Shen et al., 2023).

**Figure 1.**
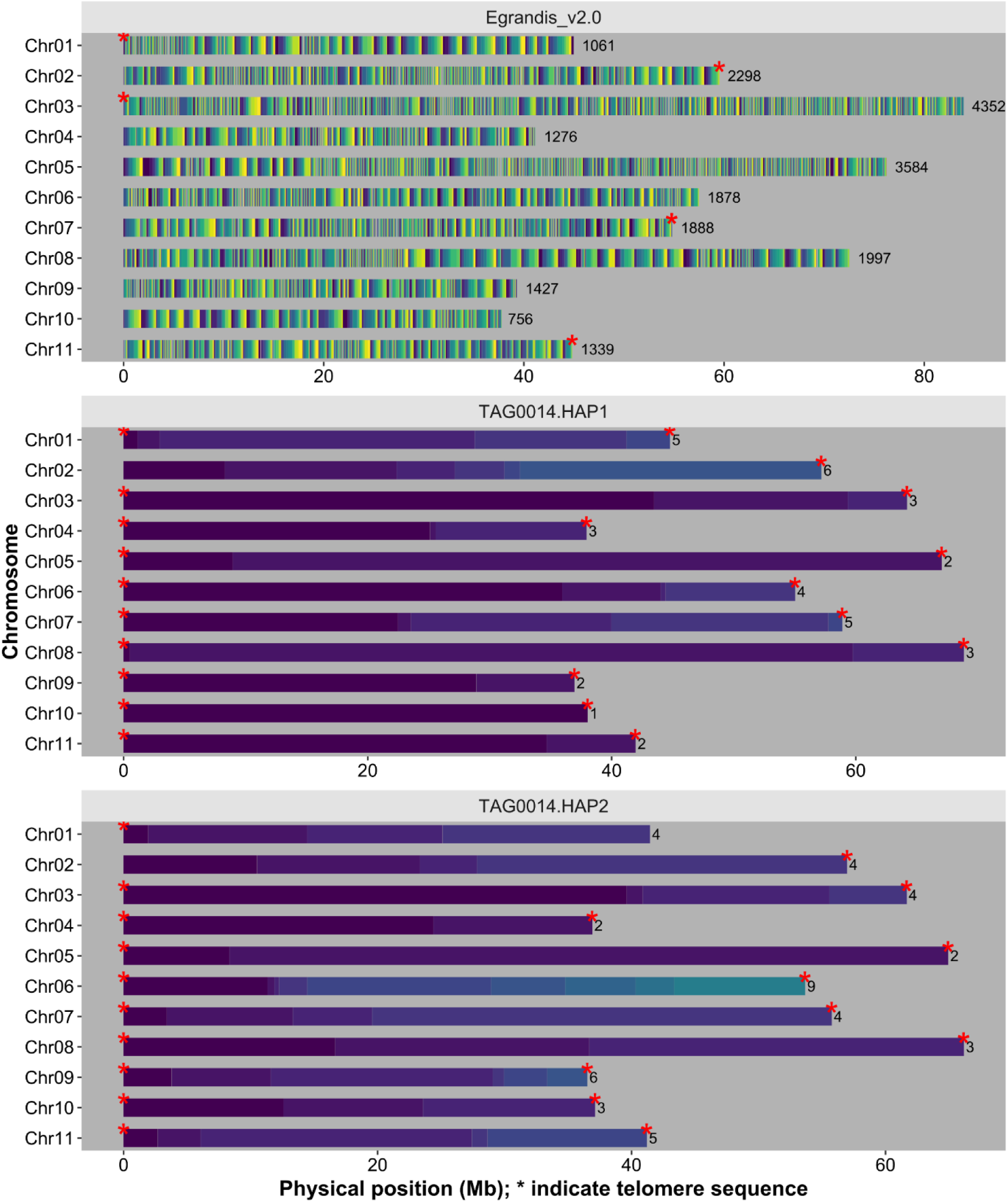
Improvement in contiguity of the *E. grandis* reference genome. Contig position of the v4.0 reference genome assemblies were generated using GENESPACE (Lovell et al., 2022). The contigs in the previous *E. grandis* v2.0 reference assembly (top panel), TAG0014 HAP1 assembly (middle panel) and TAG0014 HAP2 assembly (bottom panel) are shown as continuous blocks of a single colour. A yellow-blue palette of 20 colours was selected for all assemblies i.e. each cycle through colours represents 20 contigs. Telomere sequences are shown by a red asterisk and the numbers of contigs that make up individual chromosome scaffolds are shown on the right. Chromosome position and size are indicated on the x-axis, and chromosome number on the y-axis.

**Table 1.** Genome assembly and annotation statistics for the *E. grandis* v2.0 reference and the new phased TAG0014 (v4.0) HAP1 and HAP2 assemblies.

|  | *E. grandis* v2.0 | TAG0014 HAP1 | TAG0014 HAP2 |
| --- | --- | --- | --- |
| Assembly Size (Mbp) | 691.3 | 570.8 | 552.4 |
| Number of contigs | 32,835 | 37 | 46 |
| Contig N50 (Mbp) | 0.0672 | 28.9 | 16.7 |
| Number of scaffolds | 4,943 | 12 | 11 |
| Scaffold N50 (Mbp) | 57.5 | 57.2 | 55.8 |
| Assembly BUSCO (%) | 97.9 | 98.5 | 98.3 |
| Number of telomeres | 5 | 21 | 20 |
| Genome in scaffolds > 50 kbp (%) | 94.2 | 100 | 100 |
| Number of genes (primary transcripts) | 36,349 | 35,929 | 35,583 |
| Number of mono-exonic genes | 7,611 | 6,182 | 6,165 |
| Number of multi-exonic genes | 28,738 | 29,747 | 29,418 |
| Repeat content (%) | 44.5 | 44.99 | 43.78 |
| Number of genes with long introns (> 10 kbp) |  | 438 | 472 |
| Alternative transcripts | 9,931 | 28,647 | 28,368 |
| Annotation BUSCO (%) | 93.8 | 99.4 | 99.5 |

TAG0014 is highly heterozygous, with ∼5.2 million heterozygous SNPs and ∼534,000 heterozygous small INDELS across the two TAG0014 haplotypes (**Supplementary Table 3**), further motivating the establishment of a new reference for the species. We sequenced and assembled 570.8 Mbp and 552.4 Mbp for HAP1 and HAP2 of TAG0014. Both assemblies contained 11 chromosomes, with a contig N50 of 28.9/16.7 Mbp, and a scaffold N50 of 57.2/55.8 Mbp (**Figure 1, Table 1, Supplementary Table 1**). The haplotypes included 21/20 of the 22 telomeres supporting the improved completeness of the assembly (**Figure 1, Table 1**). Genome assembly completeness was >98.3%, with only 3.2%/2.5% duplicated complete BUSCOs, reflecting the haploid nature of the assemblies (**Supplementary Table 2**).

The HAP1/HAP2 genomes consisted of 44.99%/43.78% transposable elements (TEs, **Table 1, Supplementary Table 5**). Most repeats were unknown elements (27.51%/26.34% of the genome), followed by *Copia* LTRs (8.96%/9.18%) and *Gypsy* LTRs (3.58%/3.45%) (**Figure 3, Table 1, Supplementary Table 5**). LTR retrotransposons had an average length of 1,327/1,324 bp for HAP1/HAP2.

A combination of *ab initio* prediction, transcript evidence from PacBio Iso-Seq data of three tissue pools and short-read RNA-Seq data from 15 tissues was used to predict protein-coding regions. We annotated 35,929 and 35,583 protein-coding genes, with 28,647 and 28,368 alternative transcripts in HAP1 and HAP2, respectively (**Figure 3, Table 1, Supplementary Table 6**). We found 6,182 and 6,165 mono-exonic genes for HAP1 and HAP2, respectively (**Supplementary Table 7**). Similarly, there were 29,747 and 29,418 multi-exonic genes for HAP1 and HAP2 (**Supplementary Table 7**). The average number of exons per gene was 5.1 for HAP1 and 5.0 for HAP2 (**Supplementary Table 6**). More than 99.4% of the 1,614 embryophyta_odb10 BUSCO genes were annotated (**Supplementary Table 2**). The average genome-wide gene density was one gene every 15.89 kbp in HAP1 and one gene every 16.04 kbp in HAP2 with an average gene length of 4,020 bp (HAP1) and 3,994 bp (HAP2) and a maximum intron length of 87,554/39,127 (HAP1/HAP2) bp (**Supplementary Table 7**). A total of 438 and 472 genes contain introns larger than 10 kbp (**Table 1**). We found that a total of 1,154/1,137 genes from the v2.0 BRASUZ1 reference were merged into 544/536 genes in TAG0014 HAP1/HAP2, of which 200/189 genes were merged into 82/75 genes with long introns (> 10 kbp).

### An improved reference genome for E. grandis

The two haplotype assemblies were approximately 70 - 90 Mbp smaller than the v2.0 reference. The assemblies also contained fewer gaps (0.1%), compared to the 7.4% of gaps observed in the v2.0 reference. Overall, the two assemblies were over 700-fold more contiguous than the v2.0 reference (**Figure 1**). This is due to a combination of HiFi long read technology, much higher sequencing depth (65.5x), and the use of Omni-C contact maps which are more informative for fine-scale genome scaffolding than the genetic linkage maps used for the v2.0 assembly. Finally, the assembly includes 21 and 20 of the 22 telomeres in each haplotype, which is 4-fold more than the v2.0 reference (**Figure 1, Table 1**).

The fact that we used the same sequencing and assembly methodologies for the two haplotypes with similar accuracy, completeness, and contiguity facilitated direct comparison of the two haplotypes to assess differences in genome features that better reflect haplotype divergence in *E. grandis* than previous studies. We acknowledge that comparison of genome-wide synteny based on pairwise assembly alignments with the current v2.0 reference genome most likely reflect assembly differences rather than true genetic differences and caution against evolutionary interpretations from these results. Therefore, when making biological interpretations and conclusions regarding synteny, we only considered comparisons between the TAG0014 HAP1 and HAP2 assemblies.

We performed pairwise synteny comparisons between the *E. grandis* v2.0 and TAG0014 HAP1, TAG0014 HAP1 vs and TAG0014 HAP2, and TAG0014 HAP2 and *E. grandis* v2.0 genome assemblies. We found that 370.2/370.7 Mbp (approximately 53.6%) of the HAP1/HAP2 assemblies were syntenic to the v2.0 genome assembly. In comparison, the HAP1 and HAP2 assemblies were more syntenic to each other with 396.0/396.3 Mbp (69.4%/71.7%) being syntenic (**Figure 2, Supplementary Table 8**). This is expected for genome assemblies generated using the same sequencing and assembly methods and illustrates why caution should be applied when comparing genomes sequenced and assembled with different technologies. Overall, the *E. grandis* genomes were largely collinear with 1,448 to 1,474 translocations (total 51.0 Mbp and 55.1 Mbp) and 38 to 39 inversions larger than 10 kbp (total 18.2 Mbp and 7.6 Mbp; **Figure 2**). In comparison, the TAG0014 genomes were more collinear with 1,018 translocations (29.3 Mbp) and 22 inversions (9.25 Mbp) larger than 10 kbp between the two genomes. We observed more regions that do not align between the v2.0 reference and the HAP1 (107.7 Mbp, 15.6%) and HAP2 (114.9 Mbp, 16.6%) assemblies than between HAP1 and HAP2 (65.8 – 75.5 Mbp, 11.9 – 13.2%, **Figure 2, Supplementary Table 8**). In addition, there were fewer translocations and duplications between HAP1 and HAP2 than between them and the v2.0 reference assembly. This may reflect the difficulty of assembling and collapsing such regions into a single haplotype representation of the diploid genome of outbred organisms such as *E. grandis*. The size of SVs ranged from 100 bp to 7.96 Mbp, with the largest event detected being an inversion between HAP1 and HAP2 of TAG0014 (**Figure 2**).

**Figure 2.**
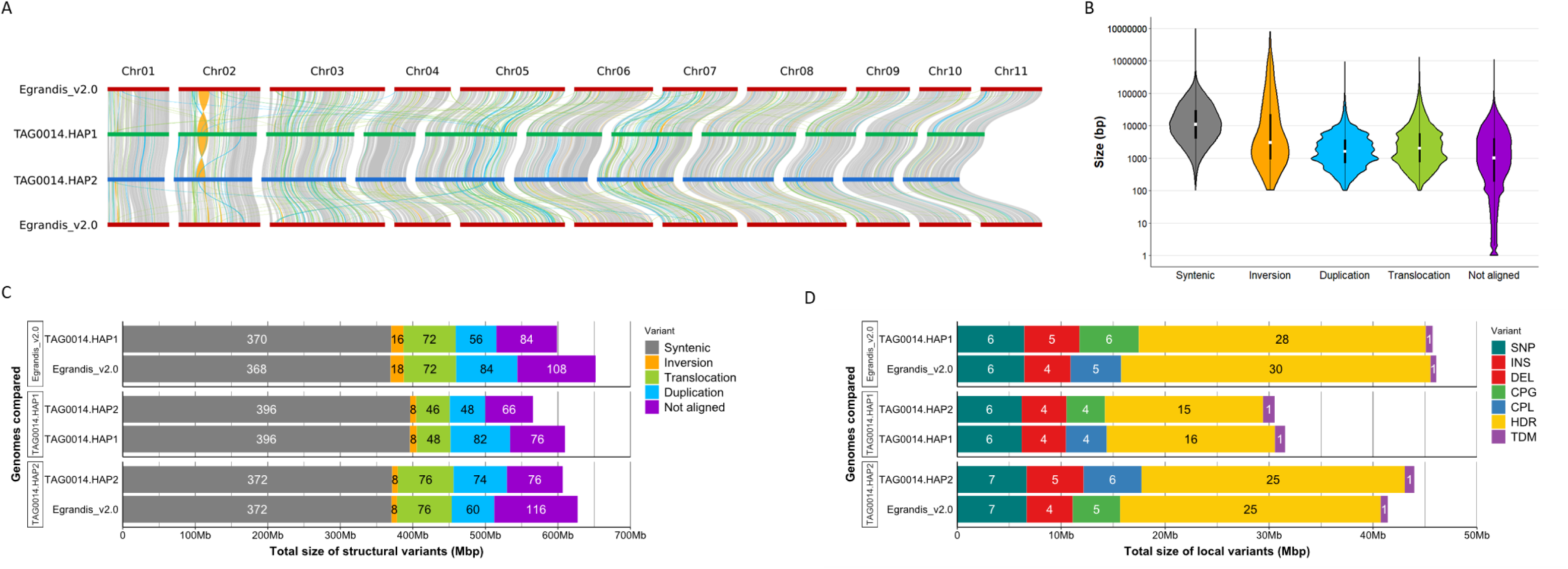
Distribution and size of structural rearrangements and local variants. **A**) Position of structural variants (SVs) across and between chromosomes of the v2.0 reference assembly (dark red), TAG0014 HAP1 (green) and TAG0014 HAP2 (blue) as calculated with SyRI (Goel et al., 2019) and visualised with plotsr (Goel & Schneeberger, 2022). Chromosome numbers are indicated at the top. SVs colours are indicated in the same colour scheme as shown in B and C. **B**) Size distribution of syntenic regions, SVs and regions that do not align. Variant type is shown on the x-axis and size in base pairs on the y-axis (on a logarithmic scale). **C**) Total size contribution of syntenic regions, SVs and non-aligned regions for each pairwise assembly comparison. The reference genome for each comparison is provided as facet labels on the y-axis, along with the specific genome compared. Total size of regions comprising each the variant type is indicated on the x-axis in megabase pairs (Mbp). **D**) Size contribution of local variants (LVs) for each pairwise assembly comparison. The reference genome for each comparison is provided as facet labels on the y-axis, along with the specific genome compared. The total size of the variant type is indicated on the x-axis in Mbp. Variants compared are SNPs, insertions (INS), deletions (DEL), copy gains (CPG), copy losses (CPL), highly diverged regions (HDR) and tandem repeats (TDM).

We observed fewer local variants (LVs, smaller than 50 bp within larger alignment blocks, including insertions, deletions, SNPs, copygains, copylosses, tandem repeats and highly diverged regions) between the HAP1 and HAP2 assemblies than between the TAG0014 assemblies and the v2.0 reference genome. For LVs, there are almost twice as many highly diverged regions (single basepairs missing in both assemblies within an alignment block) between the v2.0 reference and TAG0014 than between HAP1 and HAP2 of TAG0014. We also noted more SNPs (6.5 million vs 6.2 million), insertions (4.9 million vs 4.3 million), deletions (4.9 million vs 4.3 million), copygains (1,516 vs 1,274), copylosses (1,533 vs 1,219) and tandem repeats (320 vs 286) on average between the v2.0 reference and the TAG0014 haplotypes than between the TAG0014 haplotypes (**Figure 2, Supplementary Table 9**). The total size of LVs was larger when comparing TAG0014 haplotypes to the v2.0 reference (45.9 Mbp for HAP1 and 42.7 Mbp for HAP2) than between the TAG0014 haplotypes (31.0 Mbp, **Figure 2, Supplementary Table 9**).

The TAG0014 haplotypes had similar gene content with 128.48 Mbp (HAP1) and 126.93 Mbp (HAP2) while the gene content of the v2.0 reference was lower at 111.95 Mbp. This may also explain the lower BUSCO completion score of the v2.0 genome annotation. The repeat content of the v2.0 reference (44.5%) was similar to that of the TAG0014 assemblies (44.9% HAP1/ 43.78% HAP2, **Figure 3, Table 1, Supplementary Table 5**).

**Figure 3.**
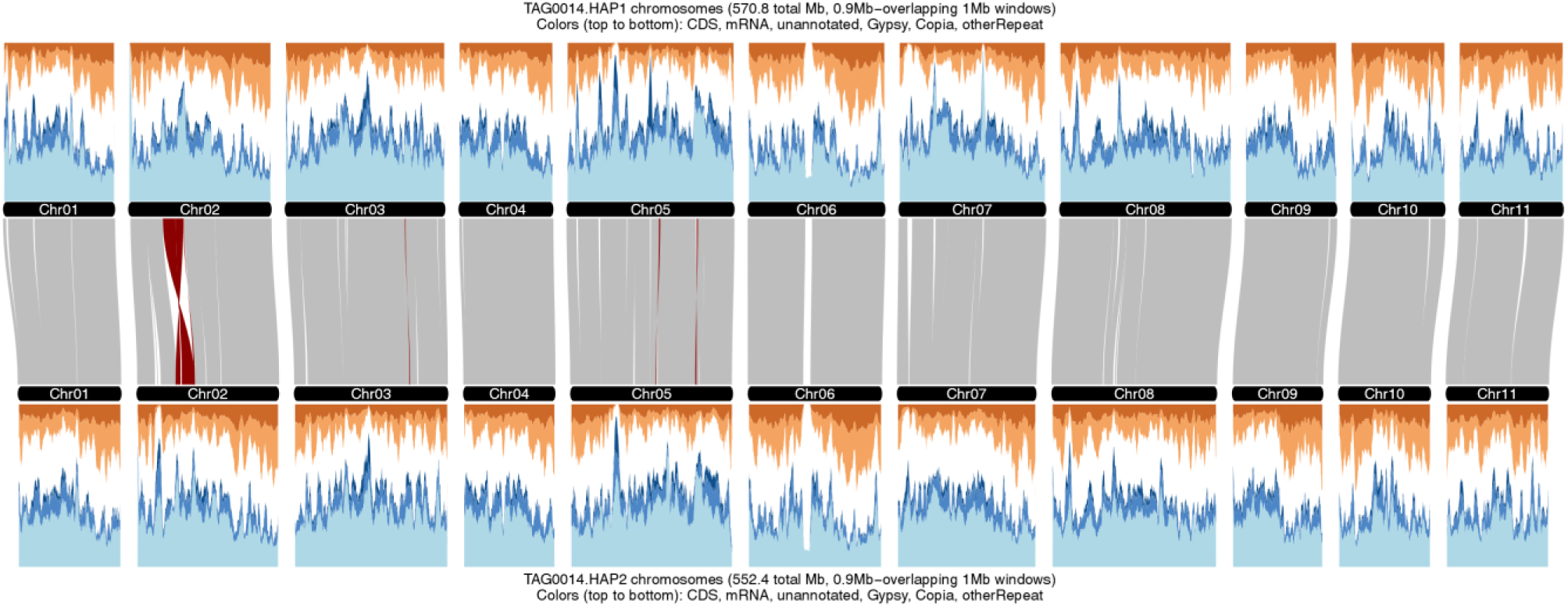
Structure and genome features of the two haplotypes of the phased TAG0014 (v4.0) genome assembly. The genome architecture between the TAG0014 HAP1 and HAP2 assemblies generated with GENESPACE. Repeat and gene density were hierarchically classified as coding sequences (CDS, dark orange), mRNA (light orange), Gypsy repeats (dark blue), Copia repeats (mid blue), other repeats (light blue) and other (white). Sliding windows of 1 Mbp are shown with 0.9 Mbp steps plotted along the horizontal axis.

Using a gene-based synteny approach, we detected high levels of collinearity between the v2.0 reference genome and the TAG0014 haplotypes and between the TAG0014 haplotypes **(Figure 3**). We found 101,490 genes in 21,149 phylogenetically hierarchical orthogroups (HOGs produced by Orthofinder), of which 16,144 were single-copy HOGs shared by all three assemblies, while 2,602 genes in 669 HOGs were assembly-specific and 6,371 genes were not assigned to a HOG.

One of the main features of the *E. grandis* v2.0 reference genome was the high number of genes in tandem duplicate arrays. Tandem duplicate arrays are still difficult to identify, but since the release of the v2.0 reference genome, new methods have been developed to define such arrays more accurately. We used GENESPACE to calculate whether the previous feature still holds true with current more accurate methods. We identified tandem arrays as described by Lovell et al., 2022 and only arrays with two or more genes were selected. We found 2,235, 2,232 and 2,106 arrays for HAP1, HAP2 and v2.0 respectively which contained 6,493, 6,346 and 5,488 genes (**Supplementary Table 10**). This was fewer than the 12,570 (34%) genes described for the v2.0 reference but still made up 18.40%, 17.83% and 15.10% of the respective total number of genes (**Supplementary Table 10**). Further analyses revealed that the v2.0 reference genome has larger arrays (with more genes in the array) than TAG0014 HAP1/HAP2 for 1,799/1,804 OGs and smaller arrays for 2,047/2,053 OGs, with 1,726 arrays larger than both haplotype-phased assemblies and 1,764 smaller than both.

## Discussion

The current collapsed diploid genome reference for *E. grandis* (v1.1: BRASUZ1; Myburg et al., 2014, later improved by Bartholome et al., 2015; v2.0) has limitations for studying large SVs that contribute to pangenome diversity in the species. Due to the technologies and methodologies used for genome sequencing and assembly, the v2.0 assembly likely contains many instances of co-assembly of partially overlapping alternative haplotypes collapsed into a pseudo-haploid reference sequence with many cases of haplotype switching in the heterozygous regions of the genome. In this context, the v2.0 reference is not representative of the genome of any *E. grandis* individual (**Figure 4**). Using a combination of PacBio HiFi, Illumina, Omni-C, RNA-Seq and PacBio Iso-Seq sequencing data, we produced an improved, haplotype-phased reference genome assembly and annotation for *E. grandis.* The phased v4.0 reference will enable studies aimed at understanding the effects that sequence and SVs have on gene regulation and local adaptation and provide access to a new source of genetic variation for molecular breeding.

**Figure 4.**
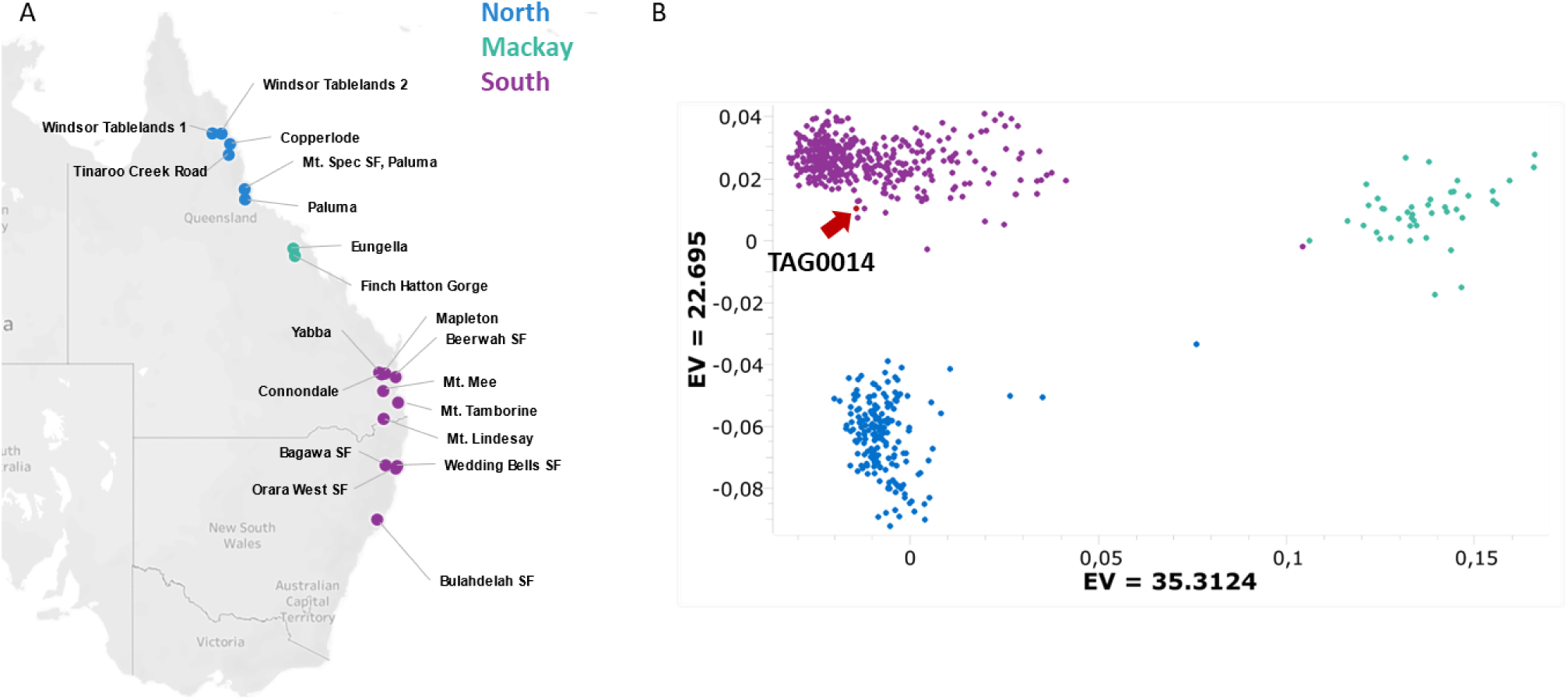
Genetic relatedness of the reference individual TAG0014 compared to the natural distribution and population diversity of *E. grandis*. (modified from Mostert-O’Neill et al., 2021). **A**) Broad geographic regions where *E. grandis* is found. Provenances are coloured and grouped by broad geographic regions, North (blue), Mackay (green) and South (purple). **B)** Population structure is depicted based on Principal Component Analysis (PCA) with TAG0014 included as shown by the red arrow based on 17,036 SNPs.

The v4.0 haplotype phased assemblies are 70 to 90 Mbp smaller than the v2.0 reference, likely reflecting that the collapsed diploid assembly is inflated relative to the two haplotype phased assemblies in the v4.0 reference. Other recent phased assemblies for *E. grandis* show a similar trend (Lotter et al., 2023; Shen et al., 2023). The TAG0014 phased assemblies are more contiguous and complete than the v2.0 reference, with almost no gaps and almost all telomeres assembled. With the aid of proximity ligation data (Omni-C), all contigs could be placed onto the eleven chromosomes of each haplotype assembly.

We find that the TAG0014 haplotypes have similar number of genes to that of the v2.0 reference (35,929/35,583 HAP1/HAP2 compared to 36,349 in v2.0) and have discovered 438 and 472 genes that have introns longer than 10 kbp. These genes merge 200/189 of the original v2.0 gene models and were only discoverable due to the use of PacBio Iso-Seq long-read RNA sequencing data. The v4.0 gene models are also more complete than the current v2.0 annotation (> 99.5% vs 93.8%) and with fewer duplicate genes (3.5%/2.8% HAP1/HAP2 vs 5.0% for v2.0, **Supplementary Table 2**). The repeat content of the TAG0014 assemblies is similar to those of the v2.0 assembly (44.99%/43.78% HAP1/HAP2 vs 44.5% for v2.0, **Table 1, Supplementary Table 5**). This suggests that the change in assembly size is not due to a general change in the repeat content, and that the repeat content is well captured. This also suggests that the smaller TAG0014 genome size is due to more accurate assembly of complex, heterozygous regions vs the need to collapse such regions in the v2.0 pseudo-haploid assembly.

One of the outstanding genomic features of the v2.0 reference was the large proportion of tandem duplicate genes, the largest observed for any sequenced plant genome at the time (Myburg et al., 2014). We have used improved methodologies to assess this feature (GENESPACE) and found that there are fewer genes in tandem arrays than previously stated (5,488 in v2.0 compared to 12,570 claimed by Myburg et al., 2014, **Supplementary Table 10**). However, even though there are fewer tandem duplicate arrays than claimed previously, our GENESPACE analysis indicated that the haplotype phased TAG0014 assemblies have more genes in tandem arrays than the v2.0 reference (approximately 1,000 more genes in tandem arrays, with 6,493, 6,346 genes contained in 2,235 and 2,232 arrays for HAP1 and HAP2, compared to 5,488 in 2,106 arrays for the v2.0 reference) which comprise 18.40% and 17.83% of the genes, respectively.

## Supplementary figures

**Supplementary Figure 1.**
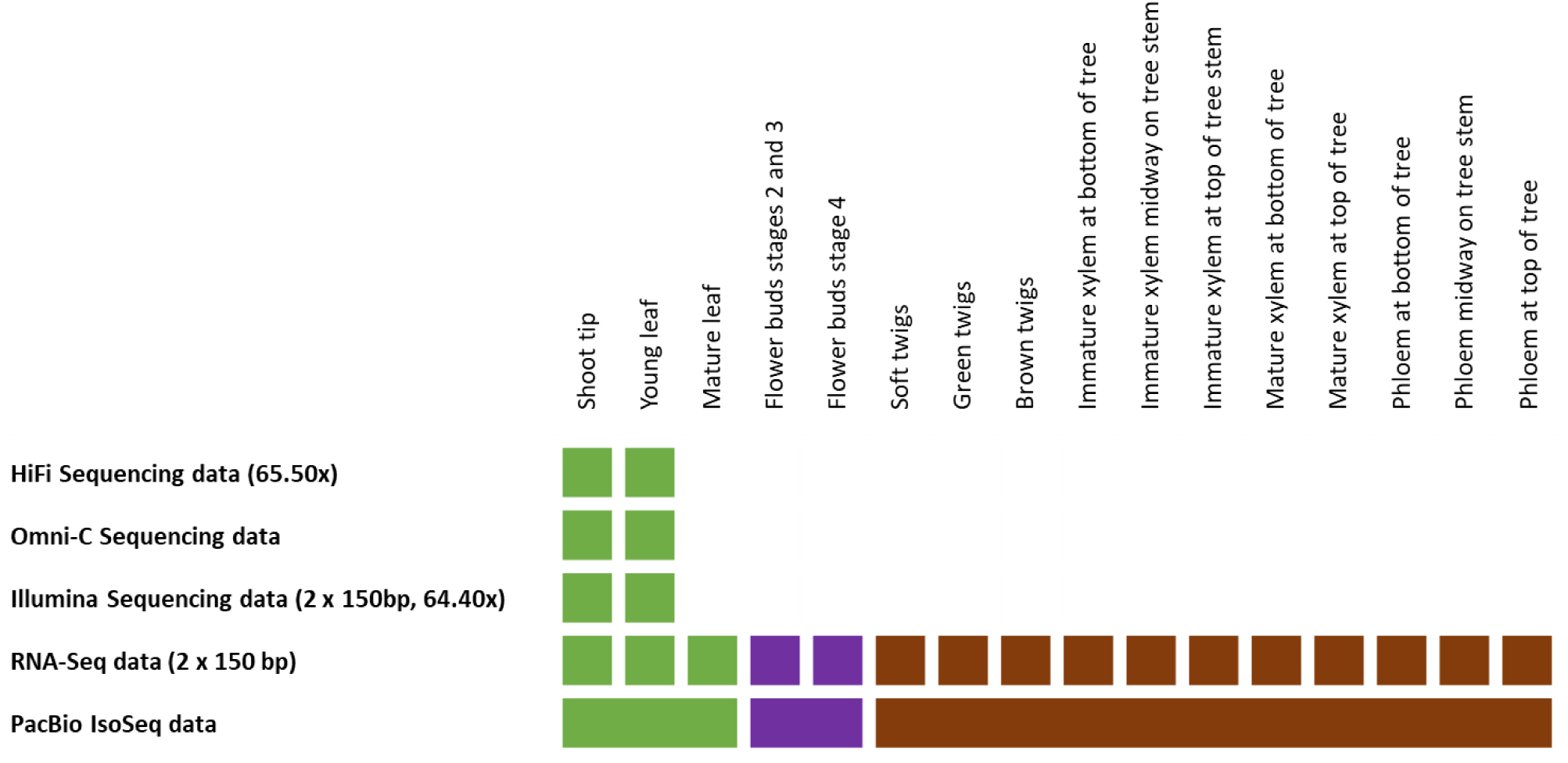
Genome resources overview. The source of the tissue used to generate the relevant sequencing data is indicated at the top. PacBio Iso-Seq sequencing data was generated in tissue pools consisting of the tissues indicated. The type of sequencing and approximate coverage is provided in brackets.

**Supplementary Figure 2.**
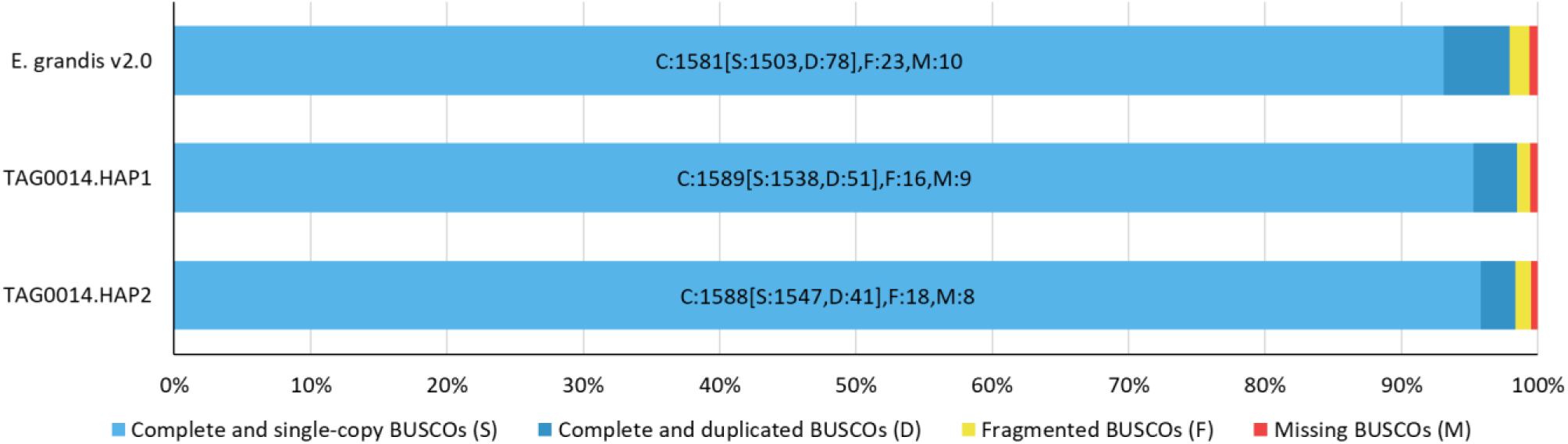
BUSCO completion scores of reference genome assemblies.

## Supplementary tables

**Supplementary Table 1.** Genome assembly and contiguity statistics.

|  | TAG0014 HAP1 | TAG0014 HAP2 |  |  |
| --- | --- | --- | --- | --- |
| QUAST assembly statistics |  |  |  |  |
| # contigs ($\geq 0$ bp) | 12 | 11 | | |
| # contigs ($\geq 1000$ bp) | 12 | 11 | | |
| # contigs ($\geq 5000$ bp) | 12 | 11 | | |
| # contigs ($\geq 10000$ bp) | 12 | 11 | | |
| # contigs ($\geq 25000$ bp) | 12 | 11 | | |
| # contigs ($\geq 50000$ bp) | 12 | 11 | | |
| Total length ($\geq 0$ bp) | 570,830,339 | 552,407,773 | | |
| Total length ($\geq 1000$ bp) | 570,830,339 | 552,407,773 | | |
| Total length ($\geq 5000$ bp) | 570,830,339 | 552,407,773 | | |
| Total length ($\geq 10000$ bp) | 570,830,339 | 552,407,773 | | |
| Total length ($\geq 25000$ bp) | 570,830,339 | 552,407,773 | | |
| Total length ($\geq 50000$ bp) | 570,830,339 | 552,407,773 | | |
| # contigs | 12 | 11 |  |  |
| Largest contig | 68,844,007 | 66,164,710 |  |  |
| Total length | 570,830,339 | 552,407,773 |  |  |
| GC (%) | 39.50 | 39.48 |  |  |
| N50 | 57,174,188 | 55,768,775 |  |  |
| N90 | 37,932,624 | 36,908,141 |  |  |
| auN | 54,538,915 | 52,725,103 |  |  |
| L50 | 5 | 5 |  |  |
| L90 | 10 | 10 |  |  |
| # N's per 100 kbp | 43.80 | 63.36 |  |  |
| Scaffolds are divided into the following sets |  |  |  |  |
| Set | Number of scaffolds | Size | Number of scaffolds | Size |
| Main genome | 12 | 570.8 Mbp | 11 | 552.4 Mbp |
| Redundant (scaffolds composed of $\geq 95\%$ 24mers $>2x$ in all scaffolds) | 63 | 8.9 Mbp | 100 | 13.9 Mbp |
| Repetitive ($\leq 250$ kbp scaffolds composed of $\geq 95\%$ 24mers $>4x$ in $\geq 5$ Mbp scaffolds) | 39 | 3.9 Mbp | 21 | 1.5 Mbp |
| Mitochondria (assembled using OatK) | 1 | 456.9 kbp | 1 | 456.9 kbp |
| Chloroplast (assembled using OatK) | 1 | 160.2 kbp | 1 | 160.2 kbp |
| Chromosome statistics |  |  |  |  |
| Scaffold | Number of contigs | Scaffold Size | Number of contigs | Scaffold Size |
| Chr01 | 5 | 44,764,794 | 4 | 41,459,349 |
| Chr02 | 6 | 57,174,188 | 4 | 56,979,195 |
| Chr03 | 3 | 64,190,686 | 4 | 61,673,418 |
| Chr04 | 3 | 37,932,624 | 2 | 36,908,141 |
| Chr05 | 2 | 67,041,112 | 2 | 64,932,641 |
| Chr06 | 4 | 55,026,902 | 9 | 53,665,386 |
| Chr07 | 5 | 58,892,594 | 4 | 55,768,775 |
| Chr08 | 3 | 68,844,007 | 3 | 66,164,710 |
| Chr09 | 2 | 36,944,981 | 6 | 36,522,489 |
| Chr10 | 1 | 38,023,239 | 3 | 37,128,649 |
| Chr11 | 2 | 41,944,707 | 5 | 41,205,020 |
| Remaining | 1 | 50,505 | 0 | 0 |

**Assembly completeness notes**
| **Total primary transcripts (34,121)** | **Number of sequences** | **Percentage of sequences** | **Number of sequences** | **Percentage of sequences** |
| --- | --- | --- | --- | --- |
| Sequences placed at 90% identity and 85% coverage | 32,663 | 95.73 | 32,577 | 95.47 |
| Sequences aligned at <50% coverage | 1,068 | 3.13 | 1,171 | 3.43 |
| Sequences not found | 390 | 1.14 | 373 | 1.09 |

**Supplementary Table 2.** BUSCO genome assembly and annotation completeness statistics for the *E. grandis* reference genomes.

| Genome | *E. grandis* v2.0 | TAG0014 HAP1 | TAG0014 HAP2 |
| --- | --- | --- | --- |
| **Assembly BUSCO** |  |  |  |
| **Percentages** | C:97.9% [S:93.1%,<br>D:4.8%], F:1.4%, M:0.7% | C:98.5% [S:95.3%,<br>D:3.2%], F:1.0%, M:0.5% | C:98.3% [S:95.8%,<br>D:2.5%], F:1.1%, M:0.6% |
| **Complete BUSCOs (C)** | 1,581 | 1,589 | 1,588 |
| **Complete and single-copy BUSCOs (S)** | 1,503 | 1,538 | 1,547 |
| **Complete and duplicated BUSCOs (D)** | 78 | 51 | 41 |
| **Fragmented BUSCOs (F)** | 23 | 16 | 18 |
| **Missing BUSCOs (M)** | 10 | 9 | 8 |
| **Total BUSCO groups searched** | 1,614 | 1,614 | 1,614 |
| **Annotation BUSCO** |  |  |  |
| **Percentages** | C:93.8% [S:88.8%,<br>D:5.0%], F:3.5%, M:2.7% | C:99.4% [S:96.2%,<br>D:3.2%], F:0.4%, M:0.2% | C:99.5% [S:96.7%,<br>D:2.8%], F:0.4%, M:0.1% |
| **Complete BUSCOs (C)** | 1,515 | 1,605 | 1,605 |
| **Complete and single-copy BUSCOs (S)** | 1,434 | 1,553 | 1,560 |
| **Complete and duplicated BUSCOs (D)** | 81 | 52 | 45 |
| **Fragmented BUSCOs (F)** | 56 | 6 | 6 |
| **Missing BUSCOs (M)** | 43 | 3 | 3 |
| **Total BUSCO groups searched** | 1,614 | 1,614 | 1,614 |

**Supplementary Table 3.** Number of heterozygous and homozygous sites fixed with polishing with Illumina (homozygous) and PacBio CCS (heterozygous) data.

| SNP/INDEL Fixing stage | TAG0014 HAP1 |  | TAG0014 HAP2 |  |
| --- | --- | --- | --- | --- |
|  | Pre SNP/INDEL fixing | Post SNP/INDEL fixing | Pre SNP/INDEL fixing | Post SNP/INDEL fixing |
| **Heterozygous SNPs** | 5,298,407 | 5,298,497 | 5,245,389 | 5,245,433 |
| **Homozygous SNPs** | 95 | 27 | 148 | 30 |
| **Heterozygous INDELs** | 534,797 | 535,451 | 533,158 | 533,785 |
| **Homozygous INDELs** | 3,804 | 51 | 4,109 | 61 |
| **Callable Bases** | 546,533,710 | 546,535,795 | 531,387,543 | 531,395,648 |

**Supplementary Table 4.** Regions that were misphased and corrected as identified by Omni-C.

| Chromosome | Haplotype | Start | End | Haplotype | Start | End |
| --- | --- | --- | --- | --- | --- | --- |
| Chr01 | HAP1 | 1,172,920 | 2,957,920 | HAP2 | 1 | 1,930,000 |
| Chr11 | HAP1 | 29,038,515 | 30,269,627 | HAP2 | 27,423,662 | 28,654,774 |

**Supplementary Table 5.**
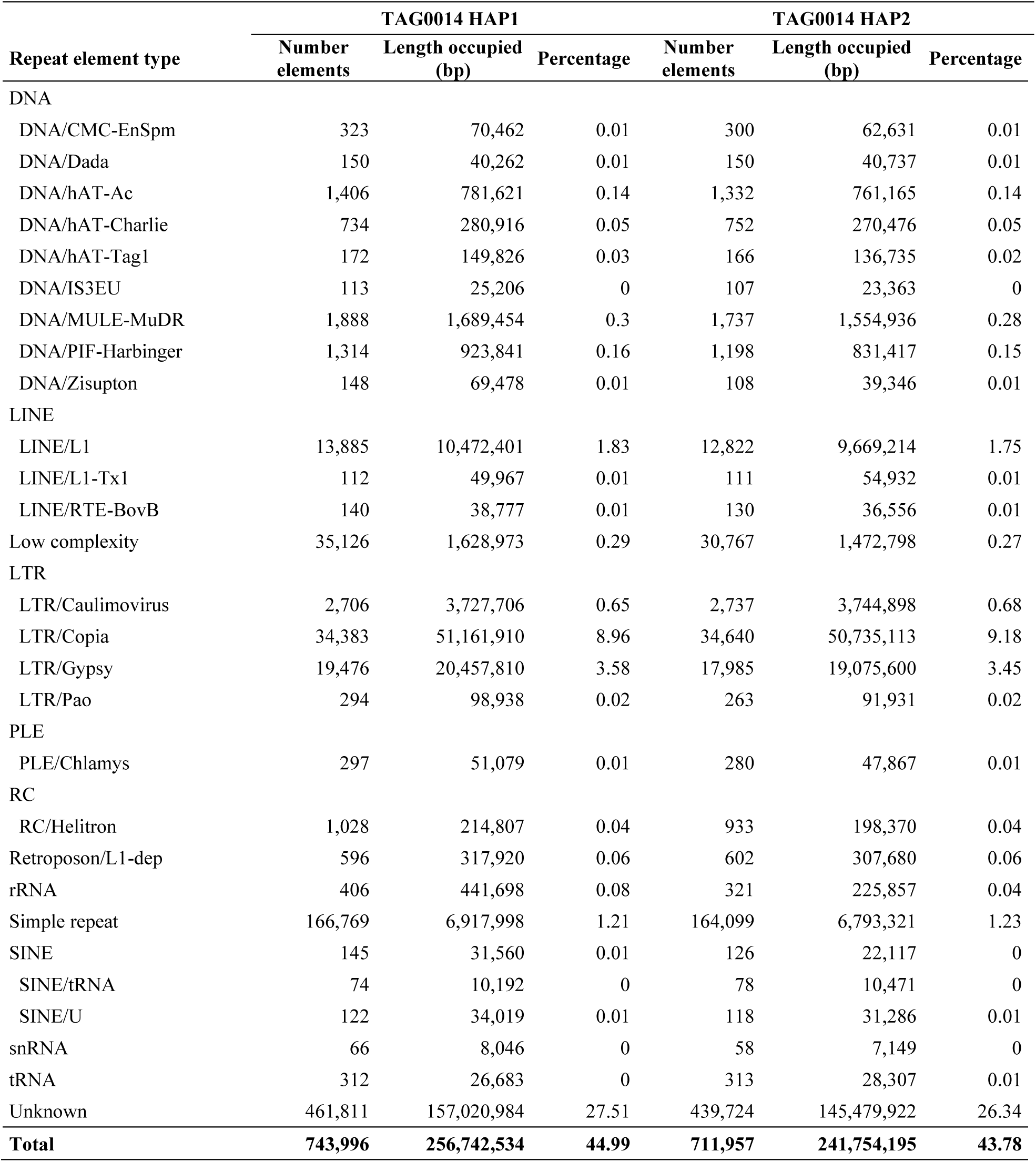
Repeat content of TAG0014 haplotype phased assemblies.

| Repeat element type | TAG0014 HAP1 |  |  | TAG0014 HAP2 |  |  |
| --- | --- | --- | --- | --- | --- | --- |
|  | Number elements | Length occupied (bp) | Percentage | Number elements | Length occupied (bp) | Percentage |
| DNA |  |  |  |  |  |  |
| DNA/CMC-EnSpm | 323 | 70,462 | 0.01 | 300 | 62,631 | 0.01 |
| DNA/Dada | 150 | 40,262 | 0.01 | 150 | 40,737 | 0.01 |
| DNA/hAT-Ac | 1,406 | 781,621 | 0.14 | 1,332 | 761,165 | 0.14 |
| DNA/hAT-Charlie | 734 | 280,916 | 0.05 | 752 | 270,476 | 0.05 |
| DNA/hAT-Tag1 | 172 | 149,826 | 0.03 | 166 | 136,735 | 0.02 |
| DNA/IS3EU | 113 | 25,206 | 0 | 107 | 23,363 | 0 |
| DNA/MULE-MuDR | 1,888 | 1,689,454 | 0.3 | 1,737 | 1,554,936 | 0.28 |
| DNA/PIF-Harbinger | 1,314 | 923,841 | 0.16 | 1,198 | 831,417 | 0.15 |
| DNA/Zisupton | 148 | 69,478 | 0.01 | 108 | 39,346 | 0.01 |
| LINE |  |  |  |  |  |  |
| LINE/L1 | 13,885 | 10,472,401 | 1.83 | 12,822 | 9,669,214 | 1.75 |
| LINE/L1-Tx1 | 112 | 49,967 | 0.01 | 111 | 54,932 | 0.01 |
| LINE/RTE-BovB | 140 | 38,777 | 0.01 | 130 | 36,556 | 0.01 |
| Low complexity | 35,126 | 1,628,973 | 0.29 | 30,767 | 1,472,798 | 0.27 |
| LTR |  |  |  |  |  |  |
| LTR/Caulimovirus | 2,706 | 3,727,706 | 0.65 | 2,737 | 3,744,898 | 0.68 |
| LTR/Copia | 34,383 | 51,161,910 | 8.96 | 34,640 | 50,735,113 | 9.18 |
| LTR/Gypsy | 19,476 | 20,457,810 | 3.58 | 17,985 | 19,075,600 | 3.45 |
| LTR/Pao | 294 | 98,938 | 0.02 | 263 | 91,931 | 0.02 |
| PLE |  |  |  |  |  |  |
| PLE/Chlamys | 297 | 51,079 | 0.01 | 280 | 47,867 | 0.01 |
| RC |  |  |  |  |  |  |
| RC/Helitron | 1,028 | 214,807 | 0.04 | 933 | 198,370 | 0.04 |
| Retroposon/L1-dep | 596 | 317,920 | 0.06 | 602 | 307,680 | 0.06 |
| rRNA | 406 | 441,698 | 0.08 | 321 | 225,857 | 0.04 |
| Simple repeat | 166,769 | 6,917,998 | 1.21 | 164,099 | 6,793,321 | 1.23 |
| SINE | 145 | 31,560 | 0.01 | 126 | 22,117 | 0 |
| SINE/tRNA | 74 | 10,192 | 0 | 78 | 10,471 | 0 |
| SINE/U | 122 | 34,019 | 0.01 | 118 | 31,286 | 0.01 |
| snRNA | 66 | 8,046 | 0 | 58 | 7,149 | 0 |
| tRNA | 312 | 26,683 | 0 | 313 | 28,307 | 0.01 |
| Unknown | 461,811 | 157,020,984 | 27.51 | 439,724 | 145,479,922 | 26.34 |
| **Total** | **743,996** | **256,742,534** | **44.99** | **711,957** | **241,754,195** | **43.78** |

**Supplementary Table 6.** Gene annotation statistics for the TAG0014 HAP1 and HAP2 reference genome assemblies.

|  | TAG0014 HAP1 | TAG0014 HAP2 |
| --- | --- | --- |
| Primary transcripts (loci) | 35,929 | 35,583 |
| Alternative transcripts | 28,647 | 28,368 |
| Total transcripts | 64,576 | 63,951 |
| For primary transcripts: |  |  |
| Average number of exons | 5.1 | 5.0 |
| Median exon length | 179 | 179 |
| Median intron length | 208 | 209 |
| **Gene model support (value is number of gene models):** |  |  |
| Any EST support | 30,914 | 30,666 |
| EST support over 100% of their lengths | 29,269 | 29,071 |
| EST support over 95% of their lengths | 29,526 | 29,315 |
| EST support over 90% of their lengths | 29,666 | 29,450 |
| EST support over 75% of their lengths | 29,954 | 29,710 |
| EST support over 50% of their lengths | 30,255 | 30,000 |
| Peptide homology coverage of 100% | 7,623 | 7,551 |
| Peptide homology coverage of over 95% | 25,444 | 25,314 |
| Peptide homology coverage of over 90% | 28,062 | 27,873 |
| Peptide homology coverage of over 75% | 31,129 | 30,789 |
| Peptide homology coverage of over 50% | 32,879 | 32,606 |
| Pfam annotation | 28,139 | 27,845 |
| Panther annotation | 32,935 | 32,563 |
| KOG annotation | 15,954 | 15,781 |
| KEGG Orthology annotation | 13,671 | 13,636 |
| E.C. number annotation | 11,333 | 11,271 |

**Supplementary Table 7.**
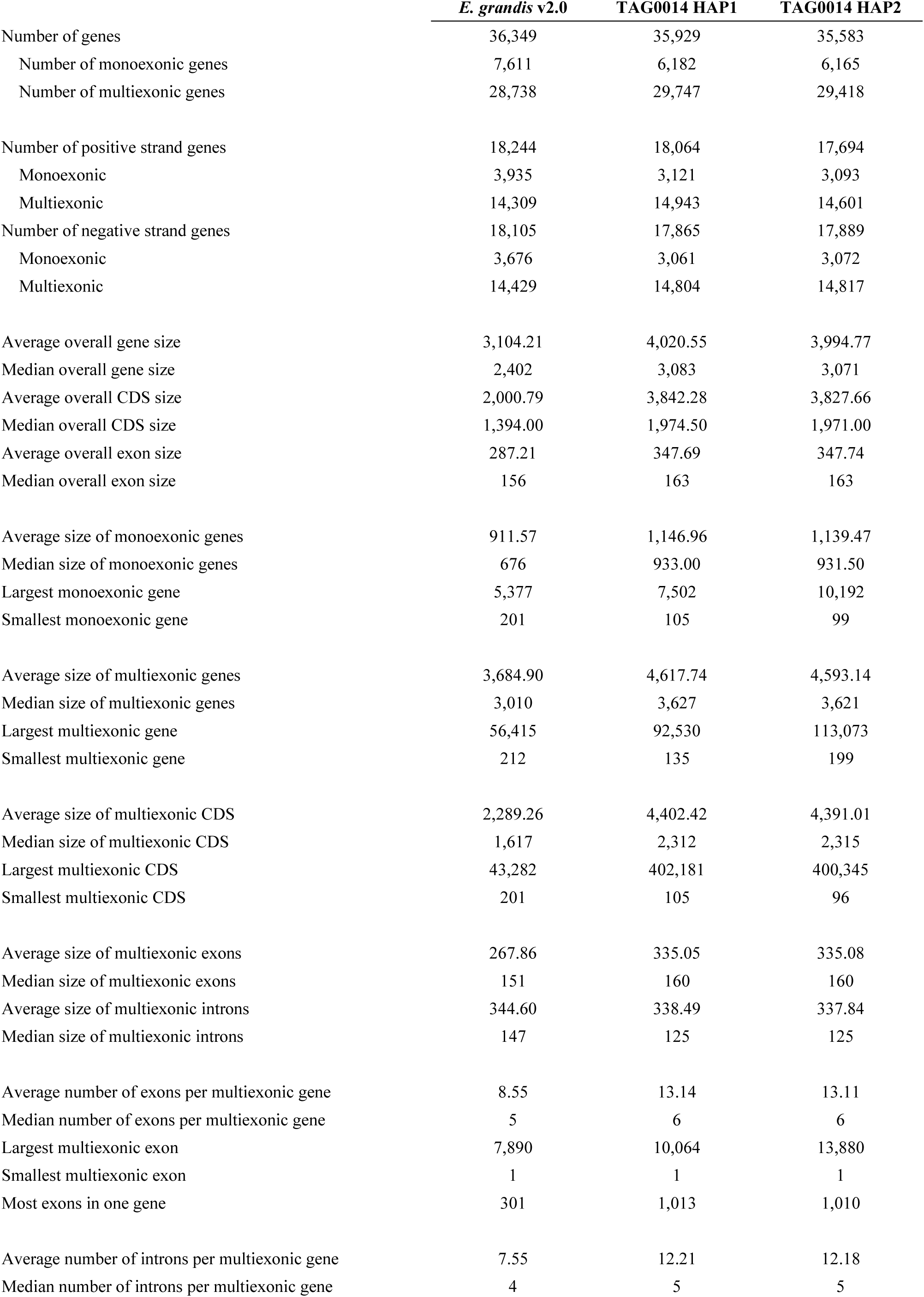
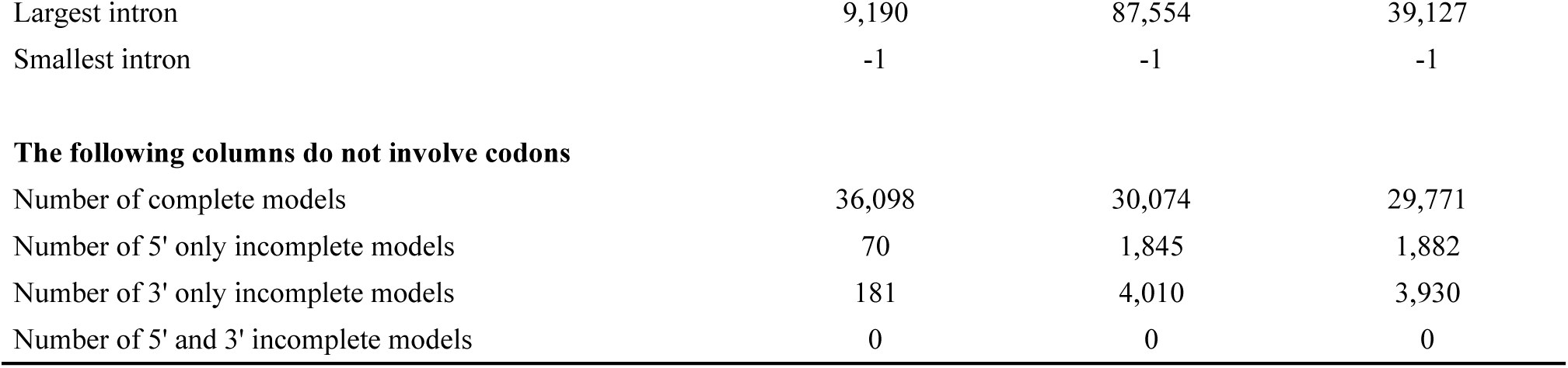
gFACS summary statistics of *E. grandis* v2.0 reference genome and TAG0014 haplotype phased assemblies.

|  | *E. grandis* v2.0 | TAG0014 HAP1 | TAG0014 HAP2 |
| --- | --- | --- | --- |
| Number of genes | 36,349 | 35,929 | 35,583 |
| Number of monoexonic genes | 7,611 | 6,182 | 6,165 |
| Number of multiexonic genes | 28,738 | 29,747 | 29,418 |
| Number of positive strand genes | 18,244 | 18,064 | 17,694 |
| Monoexonic | 3,935 | 3,121 | 3,093 |
| Multiexonic | 14,309 | 14,943 | 14,601 |
| Number of negative strand genes | 18,105 | 17,865 | 17,889 |
| Monoexonic | 3,676 | 3,061 | 3,072 |
| Multiexonic | 14,429 | 14,804 | 14,817 |
| Average overall gene size | 3,104.21 | 4,020.55 | 3,994.77 |
| Median overall gene size | 2,402 | 3,083 | 3,071 |
| Average overall CDS size | 2,000.79 | 3,842.28 | 3,827.66 |
| Median overall CDS size | 1,394.00 | 1,974.50 | 1,971.00 |
| Average overall exon size | 287.21 | 347.69 | 347.74 |
| Median overall exon size | 156 | 163 | 163 |
| Average size of monoexonic genes | 911.57 | 1,146.96 | 1,139.47 |
| Median size of monoexonic genes | 676 | 933.00 | 931.50 |
| Largest monoexonic gene | 5,377 | 7,502 | 10,192 |
| Smallest monoexonic gene | 201 | 105 | 99 |
| Average size of multiexonic genes | 3,684.90 | 4,617.74 | 4,593.14 |
| Median size of multiexonic genes | 3,010 | 3,627 | 3,621 |
| Largest multiexonic gene | 56,415 | 92,530 | 113,073 |
| Smallest multiexonic gene | 212 | 135 | 199 |
| Average size of multiexonic CDS | 2,289.26 | 4,402.42 | 4,391.01 |
| Median size of multiexonic CDS | 1,617 | 2,312 | 2,315 |
| Largest multiexonic CDS | 43,282 | 402,181 | 400,345 |
| Smallest multiexonic CDS | 201 | 105 | 96 |
| Average size of multiexonic exons | 267.86 | 335.05 | 335.08 |
| Median size of multiexonic exons | 151 | 160 | 160 |
| Average size of multiexonic introns | 344.60 | 338.49 | 337.84 |
| Median size of multiexonic introns | 147 | 125 | 125 |
| Average number of exons per multiexonic gene | 8.55 | 13.14 | 13.11 |
| Median number of exons per multiexonic gene | 5 | 6 | 6 |
| Largest multiexonic exon | 7,890 | 10,064 | 13,880 |
| Smallest multiexonic exon | 1 | 1 | 1 |
| Most exons in one gene | 301 | 1,013 | 1,010 |
| Average number of introns per multiexonic gene | 7.55 | 12.21 | 12.18 |
| Median number of introns per multiexonic gene | 4 | 5 | 5 |
| Largest intron | 9,190 | 87,554 | 39,127 |
| Smallest intron | -1 | -1 | -1 |
| **The following columns do not involve codons** |  |  |  |
| Number of complete models | 36,098 | 30,074 | 29,771 |
| Number of 5' only incomplete models | 70 | 1,845 | 1,882 |
| Number of 3' only incomplete models | 181 | 4,010 | 3,930 |
| Number of 5' and 3' incomplete models | 0 | 0 | 0 |

**Supplementary Table 8.** Summary of structural variants between the reference genomes.

| Reference genome | Query genome | Number of events | Length reference (bp) | Proportion reference (%) | Length query (bp) | Proportion query (%) |
| --- | --- | --- | --- | --- | --- | --- |
| **Syntenic regions** |  |  |  |  |  |  |
| *E. grandis* v2.0 | TAG0014 HAP1 | 14,029 | 369,254,214 | 53.41 | 370,195,762 | 64.85 |
| TAG0014 HAP1 | TAG0014 HAP2 | 13,349 | 396,000,960 | 69.37 | 396,333,150 | 71.75 |
| TAG0014 HAP2 | *E. grandis* v2.0 | 14,285 | 371,533,224 | 67.26 | 370,664,106 | 53.61 |
| **Inversions** |  |  |  |  |  |  |
| *E. grandis* v2.0 | TAG0014 HAP1 | 100 | 18,352,964 | 2.65 | 16,783,397 | 2.94 |
| TAG0014 HAP1 | TAG0014 HAP2 | 89 | 9,417,568 | 1.65 | 8,282,210 | 1.5 |
| TAG0014 HAP2 | *E. grandis* v2.0 | 114 | 7,753,224 | 1.4 | 7,569,510 | 1.09 |
| **Translocations** |  |  |  |  |  |  |
| *E. grandis* v2.0 | TAG0014 HAP1 | 9,982 | 72,043,085 | 10.42 | 72,102,959 | 12.63 |
| TAG0014 HAP1 | TAG0014 HAP2 | 7,827 | 46,538,476 | 8.15 | 46,282,393 | 8.38 |
| TAG0014 HAP2 | *E. grandis* v2.0 | 10,090 | 76,720,961 | 13.89 | 75,412,281 | 10.91 |
| **Duplications - reference** |  |  |  |  |  |  |
| *E. grandis* v2.0 | TAG0014 HAP1 | 22,713 | 84,524,332 | 12.23 |  |  |
| TAG0014 HAP1 | TAG0014 HAP2 | 21,289 | 82,095,122 | 14.38 |  |  |
| TAG0014 HAP2 | *E. grandis* v2.0 | 19,387 | 73,653,973 | 13.33 |  |  |
| **Duplications - query** |  |  |  |  |  |  |
| *E. grandis* v2.0 | TAG0014 HAP1 | 17,683 |  |  | 56,211,493 | 9.85 |
| TAG0014 HAP1 | TAG0014 HAP2 | 14,659 |  |  | 48,917,664 | 8.86 |
| TAG0014 HAP2 | *E. grandis* v2.0 | 19,526 |  |  | 58,631,124 | 8.48 |
| **Not aligned - reference** |  |  |  |  |  |  |
| *E. grandis* v2.0 | TAG0014 HAP1 | 26,566 | 107,699,067 | 15.58 |  |  |
| TAG0014 HAP1 | TAG0014 HAP2 | 20,602 | 75,546,270 | 13.23 |  |  |
| TAG0014 HAP2 | *E. grandis* v2.0 | 21,454 | 76,972,257 | 13.93 |  |  |
| **Not aligned - query** |  |  |  |  |  |  |
| *E. grandis* v2.0 | TAG0014 HAP1 | 22,602 |  |  | 83,357,649 | 14.6 |
| TAG0014 HAP1 | TAG0014 HAP2 | 18,917 |  |  | 65,828,782 | 11.92 |
| TAG0014 HAP2 | *E. grandis* v2.0 | 27,446 |  |  | 114,939,770 | 16.63 |

**Supplementary Table 9.** Summary of local variants between the reference genomes.

| Reference genome | Query genome | Number of events | Length reference (bp) | Length query (bp) |
| --- | --- | --- | --- | --- |
| **Single nucleotide polymorphisms (SNPs)** |  |  |  |  |
| *E. grandis* v2.0 | TAG0014 HAP1 | 6,453,759 | 6,453,759 | 6,453,759 |
| TAG0014 HAP1 | TAG0014 HAP2 | 6,197,893 | 6,197,893 | 6,197,893 |
| TAG0014 HAP2 | *E. grandis* v2.0 | 6,675,143 | 6,675,143 | 6,675,143 |
| **Insertions** |  |  |  |  |
| *E. grandis* v2.0 | TAG0014 HAP1 | 696,627 |  | 16,783,397 |
| TAG0014 HAP1 | TAG0014 HAP2 | 550,834 |  | 8,282,210 |
| TAG0014 HAP2 | *E. grandis* v2.0 | 626,069 |  | 7,569,510 |
| **Deletions** |  |  |  |  |
| *E. grandis* v2.0 | TAG0014 HAP1 | 494,858 | 4,448,538 |  |
| TAG0014 HAP1 | TAG0014 HAP2 | 482,009 | 4,267,853 |  |
| TAG0014 HAP2 | *E. grandis* v2.0 | 592,575 | 5,487,780 |  |
| **Copygains (CPG)** |  |  |  |  |
| *E. grandis* v2.0 | TAG0014 HAP1 | 1,626 |  | 5,703,905 |
| TAG0014 HAP1 | TAG0014 HAP2 | 1,274 |  | 3,698,389 |
| TAG0014 HAP2 | *E. grandis* v2.0 | 1,406 |  | 4,562,277 |
| **Copylosses (CPL)** |  |  |  |  |
| *E. grandis* v2.0 | TAG0014 HAP1 | 1,426 | 4,868,625 |  |
| TAG0014 HAP1 | TAG0014 HAP2 | 1,219 | 3,916,910 |  |
| TAG0014 HAP2 | *E. grandis* v2.0 | 1,639 | 5,566,419 |  |
| **Highly diverged regions (HDR)** |  |  |  |  |
| *E. grandis* v2.0 | TAG0014 HAP1 | 9,254 | 29,765,501 | 27,608,817 |
| TAG0014 HAP1 | TAG0014 HAP2 | 4,849 | 16,182,833 | 15,199,906 |
| TAG0014 HAP2 | *E. grandis* v2.0 | 9,347 | 25,316,307 | 25,061,446 |
| **Tandem repeats (TDM)** |  |  |  |  |
| *E. grandis* v2.0 | TAG0014 HAP1 | 312 | 549,240 | 659,116 |
| TAG0014 HAP1 | TAG0014 HAP2 | 286 | 970,007 | 1,121,248 |
| TAG0014 HAP2 | *E. grandis* v2.0 | 328 | 926,751 | 687,677 |

**Supplementary Table 10.** Number of tandem genes and tandem arrays in the reference genomes.

|  | *E. grandis* v2.0 | TAG0014 HAP1 | TAG0014 HAP2 |
| --- | --- | --- | --- |
| Number of genes | 5,488 | 6,493 | 6,346 |
| Number of arrays | 2,106 | 2,235 | 2,232 |
| Mean number of genes | 2.62 | 2.91 | 2.84 |
| Minimum number of genes within an array | 2 | 2 | 2 |
| Maximum number of genes within an array | 34 | 33 | 36 |
| Proportion of annotated genes (%) | 15.10 | 18.40 | 17.83 |

## Data availability

The *E. grandis* v2.0 reference is available on https://phytozome-next.jgi.doe.gov/. The TAG0014 HAP1 and HAP2 genomes will be made available on Phytozome. The raw data alongside the assembly and annotation files will be made available on NCBI under BioProject PRJNA1217046. Scripts for genome comparisons, summary statistics and to generate images are available on GitLab: https://gitlab.com/Anneri/tag0014_genome.

## Acknowledgements

Genome assemblies and sequencing data were generated by members of the Genome Sequencing Center at HudsonAlpha. All RNA-seq and Iso-Seq data QA/QC was performed by Anna Lipzen. Library construction and sequencing at the JGI was overseen by Chris Daum and Yuko Yoshinaga. The work (proposal: 10.46936/10.25585/60008105, WIP proposal ID: 508017, Myburg, Wegrzyn, Borevitz) conducted by the U.S. Department of Energy Joint Genome Institute (https://ror.org/04xm1d337), a DOE Office of Science User Facility, is supported by the Office of Science of the U.S. Department of Energy operated under Contract No. DE-AC02-05CH11231. The bioinformatics work conducted at the University of Pretoria was supported by the University of Connecticut (UConn) Computational Biology Core, who also provided computational biology support. Mondi South Africa kindly provided the plant materials used in this study.

